# Constitutive and regulatory responses of *Arabidopsis thaliana* to harmonically oscillating light

**DOI:** 10.1101/2025.02.07.637059

**Authors:** Yuxi Niu, David Fuente, Shizue Matsubara, Dušan Lazár, Ladislav Nedbal

## Abstract

The rate of net CO_2_ uptake is proportional to dim light and saturates when the light exceeds the plant assimilation capacity. This simple relationship between constant light and photosynthesis becomes intriguingly complex when the light oscillates. The rates of photosynthesis may differ between the descending and ascending phases of light oscillation. This hysteresis changes with the frequency and amplitude of the light and reports on the dynamics of the photosynthetic reactions and their regulation. Here, we investigated the chlorophyll fluorescence response of *Arabidopsis thaliana* to light oscillating with three different amplitudes: 100–200, 100–400, and 100–800 μmol photons·m^-2^·s^-1^, each with periods ranging from 1 s to 8 min. The light amplitudes and periods were chosen to represent light patterns often appearing in nature. Three genotypes were compared: wild-type Col-0 and *npq1* and *npq4* mutants that are incapacitated in the rapidly reversible energy-dependent non-photochemical quenching (qE). The experiments identified two major dynamic patterns. One was found in oscillation periods shorter than 30 s, characterized by constitutive non-linearity and hysteresis. The other was mainly formed by regulatory non-linearity and hysteresis, occurring when the oscillation periods were longer than 30 s. The mathematical model simulating the chlorophyll fluorescence dynamics qualitatively reproduced the constitutive and regulatory dynamic patterns observed in the experiments. The model simulations illustrated the dynamics of non-photochemical quencher activation, plastoquinone pool reduction, and lumen pH that form the constitutive and regulatory non-linearities. The model simulations provided mechanistic insight into molecular processes forming the plant response to oscillating light.

## 1 Introduction

The stimulus-response^1^ relationship is a fundamental concept in biology, characterizing the extent to which an organism responds to the strength, duration, or dose of a stimulus (Calabrese & Baldwin, 2003; Mattson, 2008; Pinheiro & Duffull, 2009). In plant research, this concept is commonly applied to assess optimal growth conditions or plant resistance to stress (Berry & Bjorkman, 1980; Idso & Idso, 1994; Lee *et al*., 2007; Dusenge *et al*., 2019). Another widely used stimulus-response relationship is the photosynthetic light response curve (P-I curve), which describes how the net carbon assimilation rate, P depends on the intensity of the photosynthetically active radiation, PAR or I (Evans *et al*., 1993; Ralph & Gademann, 2005; Hogewoning *et al*., 2010; Flood *et al*., 2011).

A typical P-I curve exhibits three distinct phases. Under low light intensities, the rate of photosynthesis is primarily limited by the availability of light and, therefore, it increases as a linear function of light intensity (Kiss *et al*., 2008; Krah & Logan, 2010; Murchie & Niyogi, 2011; Hasan & Cramer, 2012). Under high light, photosynthesis is limited by the electron transport and the capacity of the Calvin-Benson-Bassham cycle (Murchie & Niyogi, 2011; Hasan & Cramer, 2012; Hoh *et al*., 2023) and, therefore, increases little when PAR is increased. Thirdly, the absorbed light energy that exceeds the assimilation capacity of the plant can cause photodamage and, ultimately, photoinhibition (Krause, 1988; Aro *et al*., 1993; Allahverdiyeva & Aro, 2012) that is manifested by decreasing photosynthesis in increasing light.

The P-I curve of photosynthesis can be determined by measuring steady-state rates of net CO_2_ uptake or net O_2_ evolution in different light intensities (Evans *et al*., 1993; Ögren & Evans, 1993). An alternative way of estimating the rate of photosynthesis in relation to light intensity is by measuring the relative electron transport rate (ETR) of photosystem II (PSII). The ETR is calculated from the chlorophyll fluorescence measured by the pulse amplitude modulation (PAM) technique that probes the actinic effects of the applied actinic light, combined with saturation pulses of light that transiently close PSII reaction centers (Schreiber, 2004).

The P-I curve, measured through O_2_, CO_2_, or chlorophyll fluorescence, characterizes a steady-state photosynthesis response to constant light exposure. Such a P-I curve represents a fundamental plant stimulus-response which, however, cannot be used to understand the plant behavior in rapidly changing light that often occurs in nature (Way & Pearcy, 2012; Smith & Berry, 2013; Kaiser *et al*., 2018). The light fluctuations in different environments can be roughly classified by their typical frequencies and amplitudes (Tab.1). This is, however, only a crude characterization, and there is an endless number of light fluctuation patterns that occur in nature, each pattern potentially leading to different plant responses. On a trivial level, this variability can be illustrated by plant responses to diurnal light modulation in a square, on-off form compared with light modulation that gradually increases from morning to noon and decreases toward the evening as, e.g., in Fondy *et al*. (1989). The plant responses are different, although the period, duty cycle, and total photon energy per day may be the same in both regimes. Plants will also respond differently when light is modulated by a sine function or by a square in minutes, seconds, or shorter. The sine harmonic modulation is analogous to monochromatic light in spectroscopy or a pure musical tone. Square, triangle, or other periodic modulation patterns are analogous to polychromatic light or complex sounds because they consist of multiple harmonics represented by multiple sine functions. This originates from the uniqueness of harmonic functions of sine or cosine among all other periodic stimulus patterns (Williams, 1973; Nuij *et al*., 2006)). No other modulation pattern can be used to analyze complex periodic or even fluctuating pseudo-periodic light with the clarity of harmonic functions of sine or cosine^2^. Therefore, the stimulation of plants by harmonically modulated light is a unique probe of plant response to a particular frequency and amplitude. We use harmonically modulated light that was, in this study, sine function with periods 1 s ≤ T ≤ 8 min, i.e., of frequencies 1 Hz ≥ f ≥ 2.1·10^−3^ Hz. The respective frequencies dominate natural fluctuating light patterns represented in the bold-framed part of Tab. 1.

The dynamics of photosynthesis under harmonically oscillating light were rarely studied in the past. The pioneering work of Lam and Bungay (1986) and the position paper Lam *et al*. (1986) went largely unnoticed. Also, the independently developed line of research using harmonically modulated light (Nedbal & Březina, 2002; Nedbal *et al*., 2003; Nedbal *et al*., 2005; Matous *et al*., 2006; Berger *et al*., 2007) was seldom cited. Recently, the dynamics of photosynthesis in oscillating light and sensing in the frequency domain have become a subject of renewed interest (Shimakawa & Miyake, 2018; Samson *et al*., 2019; Jose, 2021; Nedbal & Lazár, 2021; Lazár *et al*., 2022; Niu *et al*., 2023; Niu *et al*., 2024). Lately, a mathematical model has been developed specifically to support the interpretation of mechanisms that are decisive for the stimulus-response dynamics in an oscillating light (Fuente *et al*., 2024). The model correctly predicted the dispersion, i.e., the frequency dependence of the measurable reporter signals, such as the relative chlorophyll fluorescence yield for small amplitudes of light oscillations. However, the model predictions have not yet been confronted with experiments in which the light oscillation amplitudes are large, reaching the saturation in the P-I curves. Such amplitudes often occur in nature (Tab.1) and represent a relevant scenario to study.

This led us to investigate the dependence of the photosynthetic responses to light that oscillated in a broad range of intensities from sub-saturating to saturating levels. We report on the normalized chlorophyll fluorescence yield, further ChlF(t) response of *Arabidopsis thaliana* wild-type (WT) Columbia (Col-0), and its *npq1* (Niyogi *et al*., 1998) and *npq4* mutants (Li *et al*., 2000) to light that harmonically oscillates between 100 – 200, 100 – 400, and 100 – 800 μmol photons·m^-2^·s^-1^ with periods ranging between 1 s to 8 min. The results are presented in the stimulus-response form by plotting ChlF(t) against the dynamically changing light intensity. The formal concepts of constitutive and regulatory non-linearity (Bich *et al*., 2016) were already used earlier by Nedbal and Lazár (2021) and are further developed here to classify photosynthesis’s response dynamics as constitutive and regulatory hysteresis.

As argued above, the dynamic responses of photosynthesis to harmonically oscillating light of a large amplitude differ from those observed during transients from darkness to light or reverse that are often used to probe, e.g., the activation or relaxation of non-photochemical quenching, NPQ (Nilkens *et al*., 2010; Kress & Jahns, 2017). Photosynthesis responds differently to abrupt increases and decreases in light intensity, with forward and reverse reactions happening with different rate constants. For example, the conversion of zeaxanthin to violaxanthin during NPQ relaxation is catalyzed by zeaxanthin epoxidase, whose rate constant is smaller than that of violaxanthin de-epoxidase, catalyzing the reverse reactions to convert violaxanthin to zeaxanthin during NPQ activation (Niyogi *et al*., 1997; Nilkens *et al*., 2010; Jahns & Holzwarth, 2012; Kress & Jahns, 2017). While constant-light induction and dark relaxation measurements can be used to characterize specific reactions that predominate in one of these two phases, they cannot capture the dynamic responses of photosynthesis to fluctuating light. Harmonically oscillating light provides a framework to study systemic responses in both directions as a function of frequency, offering valuable information into photosynthesis dynamics.

The experimental results were further compared with the simulations obtained by a mathematical model. The original model (Fuente *et al*., 2024) was modified here to simulate the ChlF(t) data obtained in our PAM experiments. This modified model reproduced the essential features of the experimental data and explained some of them mechanistically. The residual discrepancy between the experiment and the model simulations is used here to identify knowledge gaps, better understand photosynthesis’ regulation and operational modes in a dynamic light environment, and pave the way for future systems identification.

**Table 1.** The phenomena that result in fluctuations in photosynthetically active radiation (PAR) with different characteristic periods T and frequencies f=1/T in vegetation canopies.

| Period, $T$ | $1\text{ s} > T \geq 10\text{ ms}$ | $8\text{ min} > T \geq 1\text{ s}$ | $T \geq 8\text{ min}$ |
| --- | --- | --- | --- |
| Frequency, $f$ | $100\text{ Hz} > f \geq 1\text{ Hz}$ | $1\text{ Hz} > f \geq 2.1 \cdot 10^{-3}\text{ Hz}$ | $2.1 \cdot 10^{-3}\text{ Hz} > f$ |
| Fluctuations against a low light background in vegetation understory | Leaf flutter (Roden & Pearcy, 1993) results in very brief sun flecks (Smith & Berry, 2013). | Plant swaying (Pearcy *et al.*, 1996) and transient gaps in the canopy (Chazdon & Pearcy, 1991) cause sun flecks (Pearcy *et al.*, 1996). | Earth rotation (Pearcy *et al.*, 1996) and canopy gaps (Smith & Berry, 2013) result in long-duration sun patches (Smith & Berry, 2013). |
| Fluctuations against a medium-range background occurring within canopies | Leaf flutter (Roden & Pearcy, 1993) or plant swaying (de Langre, 2008) result in brief sun flecks (Pearcy, 1990). | Transient gaps in the upper canopy (Chazdon & Pearcy, 1991), wind-induced canopy movement (Peressotti *et al.*, 2001), and intermittent cloudiness (Knapp & Smith, 1987) give rise to complex light patterns within canopies (Way & Pearcy, 2012; Kaiser *et al.*, 2018) | Earth rotation, canopy gaps, and cloud movement result in sun and cloud patches (Smith & Berry, 2013). Diurnal changes create regular irradiance changes (Chazdon & Pearcy, 1991) |
| Fluctuations against a high-light background in outer canopies | Leaf flutter in canopies creates high-intensity sun flecks and shade flecks (Pearcy *et al.*, 1996). | Effects of plant swaying (de Langre, 2008) and intermittent cloudiness (Knapp & Smith, 1987) create sun and cloud flecks (Morales & Kaiser, 2020). | Slowly variable irradiance under overcast skies (Navrátil *et al.*, 2007), cloud movement, and diurnal changes (Morales & Kaiser, 2020). |

## 2 Materials and Methods

### 2.1 Experiments

Three genotypes of *A. thaliana* were grown in a climate chamber under light intensity of approximately 100 μmol photons·m^−2^·s^−1^, including wild-type Col-0, the *npq1* mutant that cannot convert violaxanthin into zeaxanthin (Niyogi *et al*., 1998), and the *npq4* mutant that is deficient in PsbS protein (Li *et al*., 2000). Plants were cultivated under controlled environmental conditions, with a 12-hour light / 12-hour dark diurnal regime and a day / night air temperature of 26 / 20°C. Relative air humidity was maintained at 60%. Measurements were done between 38 and 43 days after sowing.

The Dual-KLAS-NIR spectrophotometer with a 3010-DUAL leaf cuvette (Heinz Walz GmbH, Effeltrich, Germany) was used to measure the instantaneous relative chlorophyll fluorescence yield F’(t) responding to the actinic light oscillations (Klughammer & Schreiber, 2016; Schreiber & Klughammer, 2016). The data were collected every 5 ms, and 20 points were averaged to improve the signal-to-noise ratio. The resulting time resolution of 0.1 s was sufficient to capture the plant response in light oscillating with 1 Hz frequency or slower. Red actinic light (630 nm) was applied to both the abaxial and adaxial sides of the leaf. The measuring light was green (540 nm) with an intensity of 6 μmol photons·m^-2^·s^-1^, and it was applied only to the abaxial side of the leaf. The plants were collected from the climate chamber before the end of the dark photoperiod and kept in darkness until measurement. Before the oscillating light measurements, each dark-adapted plant was exposed for 10 min to constant red actinic light (630 nm) to induce photosynthesis. The intensity of this constant light was set to the average of the oscillating light that followed. Thus, plants that were later exposed to light oscillating between 100 and 200 μmol photons·m^-2^·s^-1^ were acclimated to a constant light intensity of 150 μmol photons·m^-2^·s^-1^. Those exposed to oscillations ranging from 100 to 400 μmol photons·m^-2^·s^-1^ were initially subjected to a constant light of 250 μmol photons·m^-2^·s^-1^ and, similarly, a constant light intensity of 450 μmol photons·m^-2^·s^-1^ was used for plants that were later exposed to oscillations between 100 – 800 μmol photons·m^-2^·s^-1^.

Following the induction in constant actinic light, plants were exposed to harmonically oscillating light of low, medium, and high amplitudes, as described above. The sequence of oscillating light periods was similar to Niu *et al*. (2023), consisting of eight different periods that varied continuously from 8 min to 1 s: three oscillation cycles with 8 min period, five cycles each with 4 min, 2 min, 1 min, 30 s, and 10 s periods, and finally ten cycles with 5 s and 1 s periods. The light was controlled by an 8-bit digital-to-analog converter, yielding 256 light levels to cover the amplitude range of the light intensities. This led to light changes in which discrete steps approximated the sine function: the oscillations were approximated by eight light intensity steps for 100 – 200 μmol photons·m^-2^·s^-1^, 22 light intensity steps for 100 – 400 μmol photons·m^-2^·s^-1^, and 49 light intensity steps for 100 – 800 μmol photons·m^-2^·s^-1^. The light oscillation protocol and chlorophyll fluorescence responses are illustrated in Fig. 1. Three plants of each *A. thaliana* genotype were measured in three oscillating light conditions as biological replicates.

**Figure 1.**
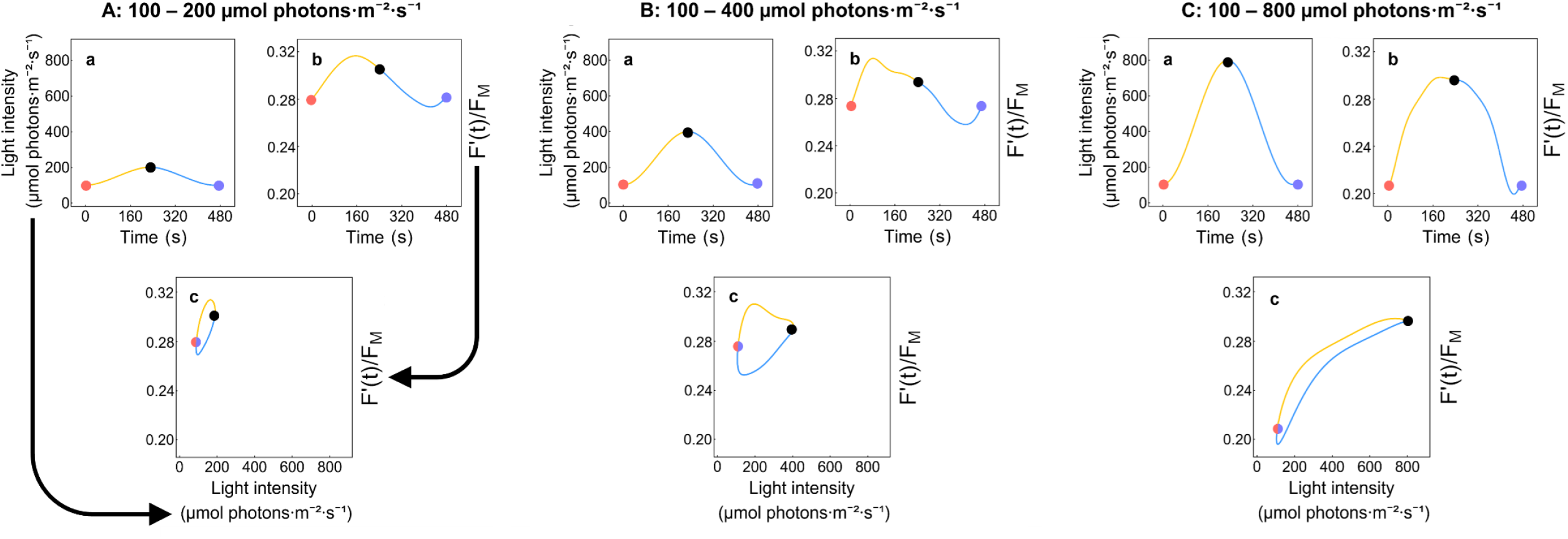
Examples of the normalized ChlF(t) = F’(t)/F_M_ of the WT strain Col-0 to light oscillating with a period T = 8 min at three different amplitudes are shown (A, B, C). The oscillation ranges were 100 – 200, 100 – 400, and 100 – 800 μmol photons·m^-2^·s^-1^. The light oscillation (Aa, Ba, Ca) starts at its minimum (red circle), continues with the ascending phase marked by the yellow line to the maximum (black circle), and concludes the period by its descending phase along the blue line to the following minimum (purple circle). The normalized ChlF(t) response is shown with the same marking in panels Ab, Bb, and Cb. The same data are shown in the stimulus-response format in Ac, Bc, and Cc, where the line and marker colors are the same as in the a and b panels. The overlapping light minima are marked by half-purple, half-red circles in c.

The dynamic patterns of ChlF(t) signals sometimes change during the first one or two cycles following the change of the oscillation period. Therefore, only the steady-state dynamic patterns^3^ that emerged in later cycles were used for the analysis. Specifically, the first cycle of the T = 8 min oscillation and the first two cycles of the other oscillation periods were excluded from the analysis to minimize aperiodic transient components.

The steady-state ChlF(t) dynamic patterns were then averaged and fitted by the function in Eq. 1 as previously done (Nedbal & Lazár, 2021; Niu *et al*., 2023).

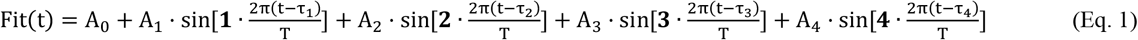

The least-square fitting was done by Microsoft Excel, yielding the offset A_0_ as well as the amplitudes (A_1,_ A_2,_ A_3,_ A_4_) and the phase shifts (τ_1_/T, τ_2_/T, τ_3_/T, τ_4_/T) of up to four harmonic components. These nine parameters characterizing separately each of the three biological replicates were averaged and statistical errors were calculated. The values of the offset A_0_ and the amplitudes of the fundamental (A_1_) as well as of the three upper harmonics (A_2,_ A_3,_ A_4_) of ChlF(t) signals under different frequencies of oscillating light are presented in Fig. SI-6. The measured raw data (Figs. SI-1, SI-2, SI-4), the fitted ChlF(t) signals (Figs. 2 to 4; Figs. SI-3 and SI-5), and also the modeled signals (Fig. 4) have been normalized by the maximum fluorescence in the dark-adapted state, F_M_: ChlF(t) =F’(t)/F_M_.

**Figure 2.**
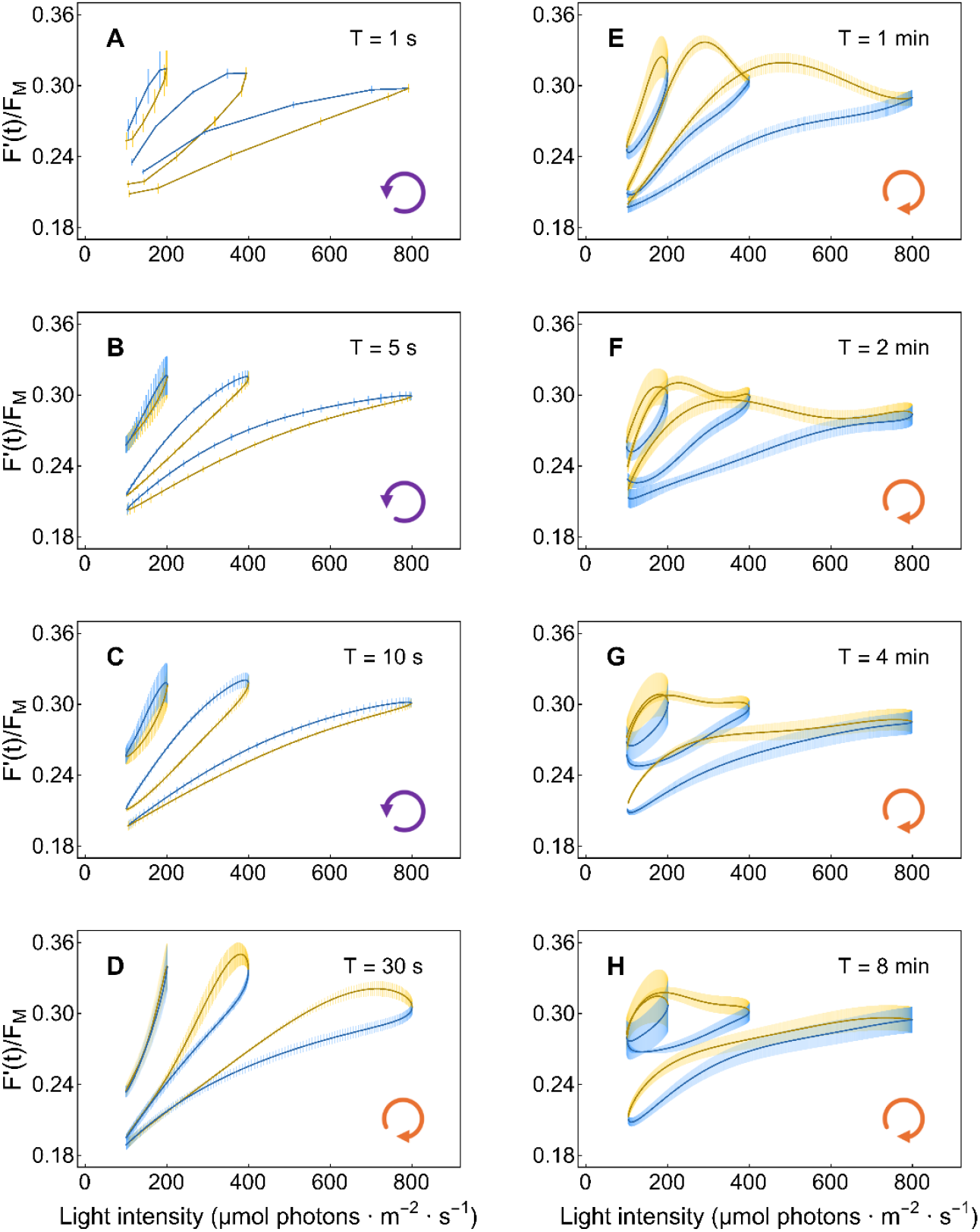
The ChlF(t)= F’(t)/F_M_ dynamics in the WT *A. thaliana* Col-0. The steady-state dynamics of ChlF(t) is shown as a function of light intensity, PAR. The light oscillated with eight different periods and three amplitudes (three different yellow-blue loops in each of the eight panels). The dynamics represent the means of three independent biological replicates, with error bars indicating standard errors (n = 3). The oscillation periods are noted in the legend of each panel, while the oscillating light intensity range for each loop is seen in the abscissa axis. The yellow-blue color code is the same as in Fig. 1, and the loop arrows at the bottom right corner of each panel indicate the orientation of the loops.

### 2.2 Mathematical model

The mathematical model BDM that was described in detail by Fuente *et al*. (2024) was further developed here to simulate photosynthetic dynamics under harmonically oscillatory light. The model (see the scheme in Fig. SI-7) consists of five ordinary differential equations representing the redox state of the PQ pool by the PQ(t) variable, the redox state of the photosystem I donors by the PI_ox_(t) variable, the lumen pH by the H_L_(t) variable, the ATP concentration by the ATP(t) variable, and a pH-dependent NPQ by the FQact(t) variable, all functions of time during the respective light oscillation. To compare model simulations with experiments, we normalize the instantaneous chlorophyll fluorescence signal measured by the PAM method, F’(t), to the maximum F_M_ attained in a multiple turnover saturating flash in a dark-adapted plant. In contrast to Fuente *et al*. (2024), we also include here the expression for the minimal chlorophyll fluorescence yield in the light-adapted state, F_0_′(t), derived by Oxborough and Baker (1997). With these modifications, one obtains the normalized instantaneous chlorophyll fluorescence yield ChlF(t):

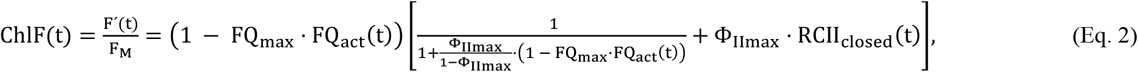

RCII_closed_(t) is a dependent variable in the BDM model representing the fraction of the closed reaction centers of PSII, and FQ_act_(t) is one of the five independent variables of BDM that represents activated quencher. FQ_max_ is a BDM parameter corresponding to the maximal NPQ when the quencher is fully activated. Φ_IImax_ (= F_V_/F_M_) is the maximum quantum yield of PSII photochemistry determined in the dark-adapted state, where F_V_ = F_M_ – F_0_ is the variable chlorophyll fluorescence yield in the dark-adapted state (reviewed in Lazár *et al*. (2005)). The details of the equation derivation are provided in the Supporting information.

## 3 Results

### 3.1 The ChlF(t) dynamics in the WT *A. thaliana* Col-0

The experimental results obtained with the WT *A. thaliana* Col-0 are represented in Fig. 2. The plots were obtained by calculating A_0_, A_1,_ A_2,_ A_3,_ A_4,_ and τ_1_/T, τ_2_/T, τ_3_/T, τ_4_/T in Eq. 1 for each experimental dataset and by calculating the corresponding averages and standard errors from measurements on three plants. The details of raw data, averages, and standard errors are shown in Fig. SI-1. The individual panels in Fig. 2 represent the normalized steady-state dynamic ChlF(t) response pattern of the plants exposed to actinic light oscillating with periods T = 1 s (A), 5 s (B), 10 s (C), 30 s (D), 1 min (E), 2 min (F), 4 min (G), and 8 min (H). Each panel shows three curves corresponding to light oscillating between 100 – 200, 100 – 400, and 100 – 800 μmol photons·m^-2^·s^-1^ of PAR. As in Fig. 1, each plotted curve consists of two phases: the first one when the light intensity oscillation starts at 100 μmol photons·m^-2^·s^-1^ and gradually increases (ascending phase, yellow line) up to the maximum value (200, 400, or 800 μmol photons·m^-2^·s^-1^) and the second one when the light intensity decreases back to 100 μmol photons·m^-2^·s^-1^ (descending phase, blue line). The orientation of the loop is further emphasized by the purple, counter-clockwise, and orange, clockwise arrows in Fig. 2. The results demonstrate that ChlF(t) responses strongly depend on the oscillating light’s period and amplitude.

The signal loops in Fig. 2 mostly show two ChlF(t) values for the same incident light intensity, depending on the illumination history; such behavior is called hysteresis. The ChlF(t) response lagged the oscillating light when the periods were shorter than 30 s, i.e., ChlF(t) was lower in the ascending light oscillation phase than in the descending phase for the same light intensity (A-C in Fig. 2). Such a delayed response is observed in many biological, chemical, and physical systems (Mayergoyz, 2003; Strogatz, 2018). We shall show further that this type of behavior is inherently caused by the constitutive non-linearities (Nedbal & Lazár, 2021) of primary photosynthetic processes and, therefore, we call it constitutive hysteresis. The loops representing the constitutive hysteresis in Fig. 2A-C were nearly elliptical for the low- and medium-amplitude oscillations (100 – 200 and 100 – 400 µmol photons·m^-2^·s^-1^) and exhibited signs of saturation in the high-amplitude oscillations (100 – 800 µmol photons·m^-2^·s^-1^).

The ChlF(t) dynamics changed strikingly when the oscillation periods increased from T = 10 s to 30 s (Fig. 2D). The loop directions changed from the counter-clockwise, which were observed with shorter periods (Fig. 2A-C), to clockwise orientation in the medium- and high-amplitude oscillations of the 30-s period (Fig. 2D). This dynamic feature was observed in all light oscillation amplitudes also with periods longer than 30 s (Fig. 2E-H). In this case, ChlF(t) was higher in the light ascending phase than in the descending phase, and ChlF(t) started to decrease already in the light ascending phase, i.e., the ChlF(t) maxima were reached before the light intensity peaked. This is caused by regulatory non-linearities, namely, by regulation lagging the light oscillation (Niu *et al*., 2023). This dynamic feature can, therefore, be called regulatory hysteresis. In our experiments with the high-amplitude light oscillations, the regulatory hysteresis dominated in the period T = 1 min and decreased as the period further increased (Fig. 2G, H). Thus, the ChlF(t) dynamics under high amplitude and slow oscillations converged to the typical steady-state P-I curves, in which the hysteresis is negligible because the photosynthesis apparatus has enough time to settle to the dynamic homeostasis for each light level and little effect of the light history is therefore expected. Interestingly, significant hysteresis remained even in the long periods (Fig. 2G, H) when the low and medium oscillation amplitudes were applied. This may indicate that the mechanisms reducing hysteresis in the high-amplitude oscillations and, therefore, the convergence to the steady state P-I requires high light. The shape of the dynamic pattern in the high-light oscillations also supports this hypothesis.

### 3.2 The ChlF(t) dynamics in the *A. thaliana npq1* and *npq4* mutants

The *npq4* mutant, which does not have the PsbS protein (Li *et al*., 2000), and the *npq1* mutant, which cannot convert violaxanthin into zeaxanthin (Niyogi *et al*., 1998), both exhibit ChlF(t) dynamic responses that differ from those of the Col-0 WT (Fig. 3). The stimulus-response plots in Fig. 3 show in the top row panels A, B, and C the differences for the period T = 1 s that was in Fig. 2 associated with constitutive hysteresis. The dynamic patterns in the two lower panel rows, T = 1 min and 4 min were dominantly formed by the regulatory hysteresis. The raw data and further details are shown in Figs. SI-3 and SI-5.

**Figure 3.**
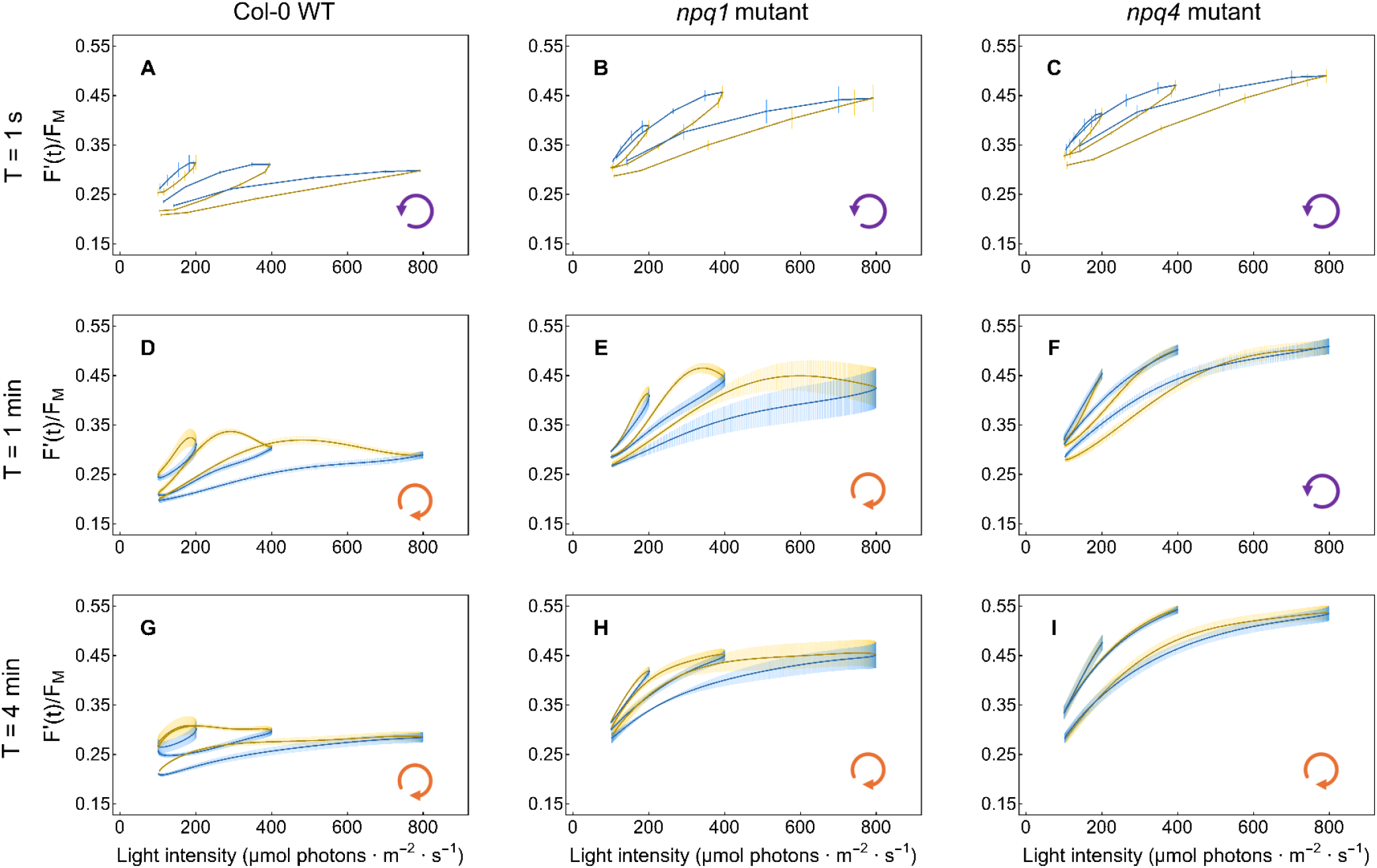
The dynamics of ChlF(t)= F’(t)/F_M_ in the *A. thaliana* Col-0 WT (left panels A, D, G), *npq1* mutant (central panels B, E, H), and *npq4* mutant (right panels C, F, I). The top row panels (A, B, C) represent ChlF(t) dynamic patterns obtained with light oscillating with period T = 1 s, the middle row (D, E, F) with T = 1 min, and the bottom row (G, H, I) with T = 4 min. The yellow-blue color code is the same as in Fig. 1. The loop arrows at the bottom right corner of each panel indicate the orientations of the loops.

The light oscillations with the short period of T = 1 s (Fig. 3A-C) elicited ChlF(t) responses that were qualitatively similar in all three genotypes except the average ChlF(t) levels, which were lower in the WT compared to the mutants. The WT plants were responding by NPQ that was partially incapacitated in the mutants, and, therefore, their average ChlF(t) yield was higher than that in the WT. As NPQ responded to the average light levels, the ChlF(t) in WT plants was lowest in high-amplitude oscillations. The opposite was the case for low-amplitude oscillations. Except for this difference in the average ChlF(t) levels, the steady-state dynamic patterns found in rapidly oscillating light in the WT and mutant plants were similar. This suggests that the T = 1 s pattern shape is formed by constitutive rather than regulatory hysteresis.

In contrast to T = 1 s, the regulatory hysteresis essentially formed the patterns found in the oscillations T = 1 and 4 min in the WT (Fig. 3D, G) and in the *npq1* mutant (Fig. 3E, H), a feature that was largely absent in the *npq4* mutant (Fig. 3F, I). The experiment shows that regulatory hysteresis occurs due to the PsbS-dependent qE that is active in the WT and *npq1*, but not in *npq4* plants. Regulatory hysteresis was weaker in T = 4 min than in T = 1 min, presumably because ChlF(t) was already approaching steady-state in the long oscillation period, and, thus, the relative effects of regulation delays were less apparent with T = 4 min than with T = 1 min. The absence of zeaxanthin-dependent qE in the *npq1* mutant and PsbS-dependent qE in the *npq4* mutant led to the amplitude of the ChlF(t) patterns in Fig. 3E, H, and Fig. 3F, I that were higher than in Fig. 3D, G that represent the WT. This shows that both types of qE are required for dynamic homeostasis, which decides the stable levels of NPQ.

Overall, ChlF(t) in WT plants was much less sensitive to light oscillations than in the mutants: The stimulus-response patterns of the WT were nearly flat even when the oscillations reached high light levels (Fig. 3G). This indicates that in the WT plants, the efficient pH-dependent qE quenching balanced the light fluctuations and responded to the mean irradiance, enabling the system to maintain energetic homeostasis despite large changes in light intensity.

### 3.3 Comparing the ChlF(t) dynamics observed in experiments with model simulations

The mathematical model, named the basic DREAM model (BDM) in (Fuente *et al*., 2024) is used and further developed here. The model scheme is repeated here in Fig. SI-7 for reading convenience. The coherence between the experiment and the model is achieved by normalizing the instantaneous chlorophyll fluorescence yield F’(t) by F_M_ both in the experiments and in the simulations (Eq. 2). Further, the model newly includes, in F’(t)/F_M_ (Eq. 2), also F_0_′(t)/F_M_, adopted from Oxborough and Baker (1997).

The experimentally measured ChlF(t) responses in the left column of Fig. 4 panels are compared with responses simulated by the model in the right column of Fig. 4. The comparison is done for short, T = 1 s and for long, 4 min periods of the light oscillations that were found above to represent the constitute and regulatory types of response hysteresis, respectively.

**Figure 4.**
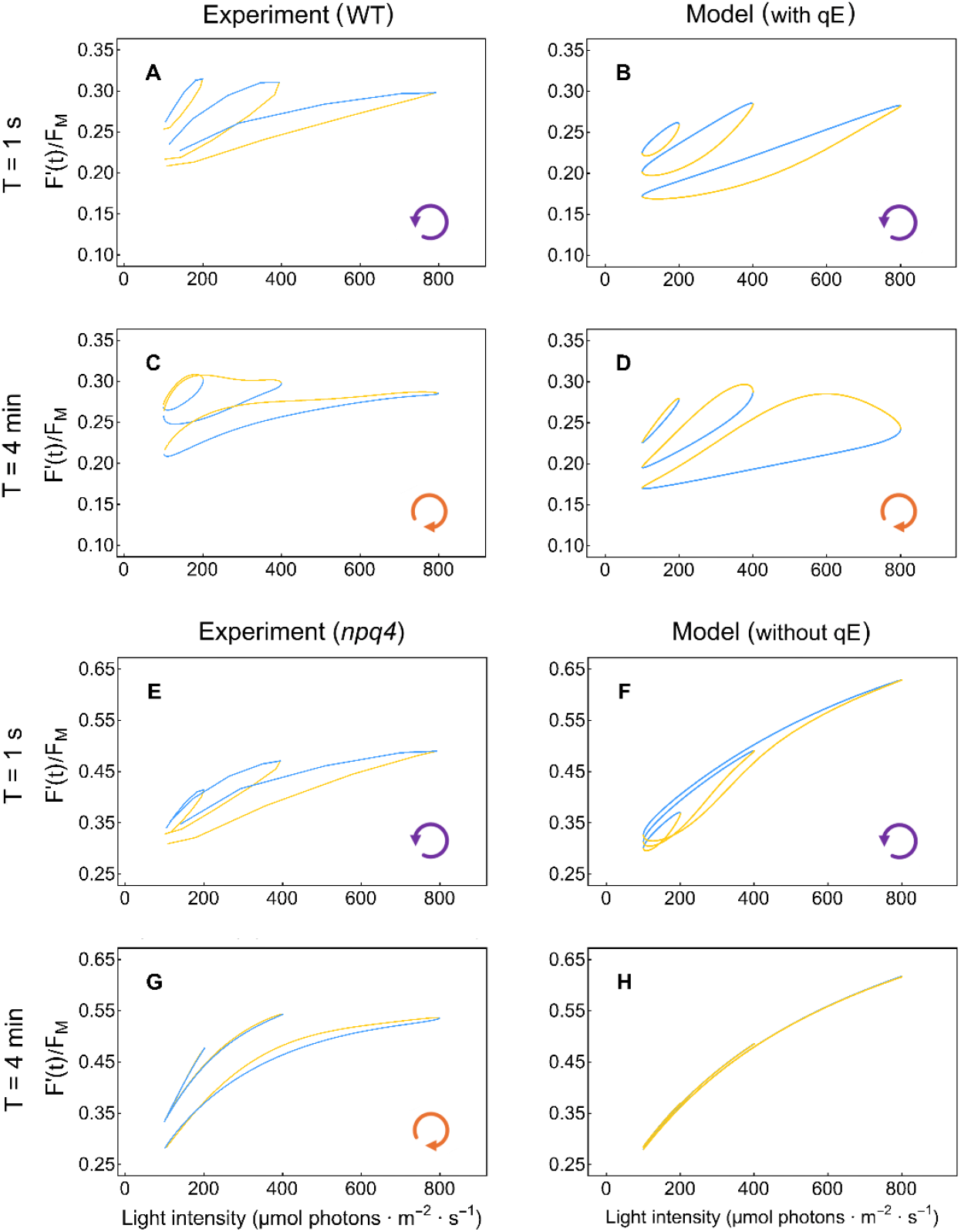
The experimentally measured ChlF(t) dynamics of WT *A. thaliana* Col-0 (panels A and C) and of the *npq4* mutant (panels E and G) are compared with simulations obtained with the modified BDM model in which qE mechanism was included (panels B and D) and in which the qE variable was set to zero (panels F and H). The data and simulations were obtained with the short and long light oscillation periods that were earlier shown to lead to the constitutive (T = 1 s) and regulatory hysteresis (T = 4 min). The color code and symbols are the same as in the previous figures.

First, the experimentally measured ChlF(t) patterns of the WT (Fig. 4A, C) are compared to the simulations by the model in which qE is included in the calculations (Fig. 4B, D). The model agrees with the T = 1 s experiment by exhibiting a significant constitutive hysteresis represented by the counter-clockwise orientation of the stimulus-response loops (cf. Fig. 4A, B). The agreement between the experiment and the model is also confirmed in T = 4 min, where the regulatory hysteresis led to the clockwise orientation of the stimulus-response loops (cf. Fig. 4C, D).

The BDM model in (Fuente *et al*., 2024) is highly reduced and lumps different mechanisms of qE in one model variable. Setting this model variable to zero, one can model the scenario without qE, as shown in Fig. 4F and 4H. The ChlF(t) patterns found in the experiments with *npq4* mutant (Fig. 4E, G) are compared with the respective patterns simulated by the model without qE (Fig. 4F, H). The *npq1* mutant is not included in the Fig. 4 comparison because it can respond by the PsbS-facilitated qE alone which is not modeled by BDM.

The ChlF(t) stimulus-response loops measured and simulated with T = 1 s reveal both the constitutive hysteresis of counter-clockwise orientation in the absence of qE (cf. Fig. 4E, F). This again confirms that the character of this dynamic feature is constitutive and independent of regulation. The stimulus-response patterns simulated with T = 4 min in Fig. 4H are void of any hysteresis. This is in line with constitutive hysteresis being negligible when the oscillation periods are much longer than the delays by the photosynthetic electron transport in filling and emptying, in the case of the BDM model, of the plastoquinone pool. The experimental data obtained for T = 4 min with the *npq4* mutant in Fig. 4G shows a similar pattern to the simulation in Fig. 4H except for a tiny regulatory hysteresis that was found with the highest oscillation amplitude.

The agreement between the experimental data and the model simulations described above is qualitative. There are also apparent differences, particularly in the ChlF(t) response ranges and in the shapes of the stimulus-response loops. These differences will be discussed further.

### 3.4 Molecular mechanisms shaping the ChlF(t) responses to oscillating light

The dynamics of three independent model variables, FQact(t), PQ(t), -log_10_(H_L_(t)) elicited by light oscillations with the periods T = 1 s and T = 4 min are simulated in Fig. 5 to interpret the molecular mechanisms that form the constitutive (T = 1 s) and regulatory hysteresis (T = 4 min). The model simulations were done while including qE in the left column of Fig. 5 and with the variable FQact(t) = 0 in the right column of Fig. 5.

**Figure 5.**
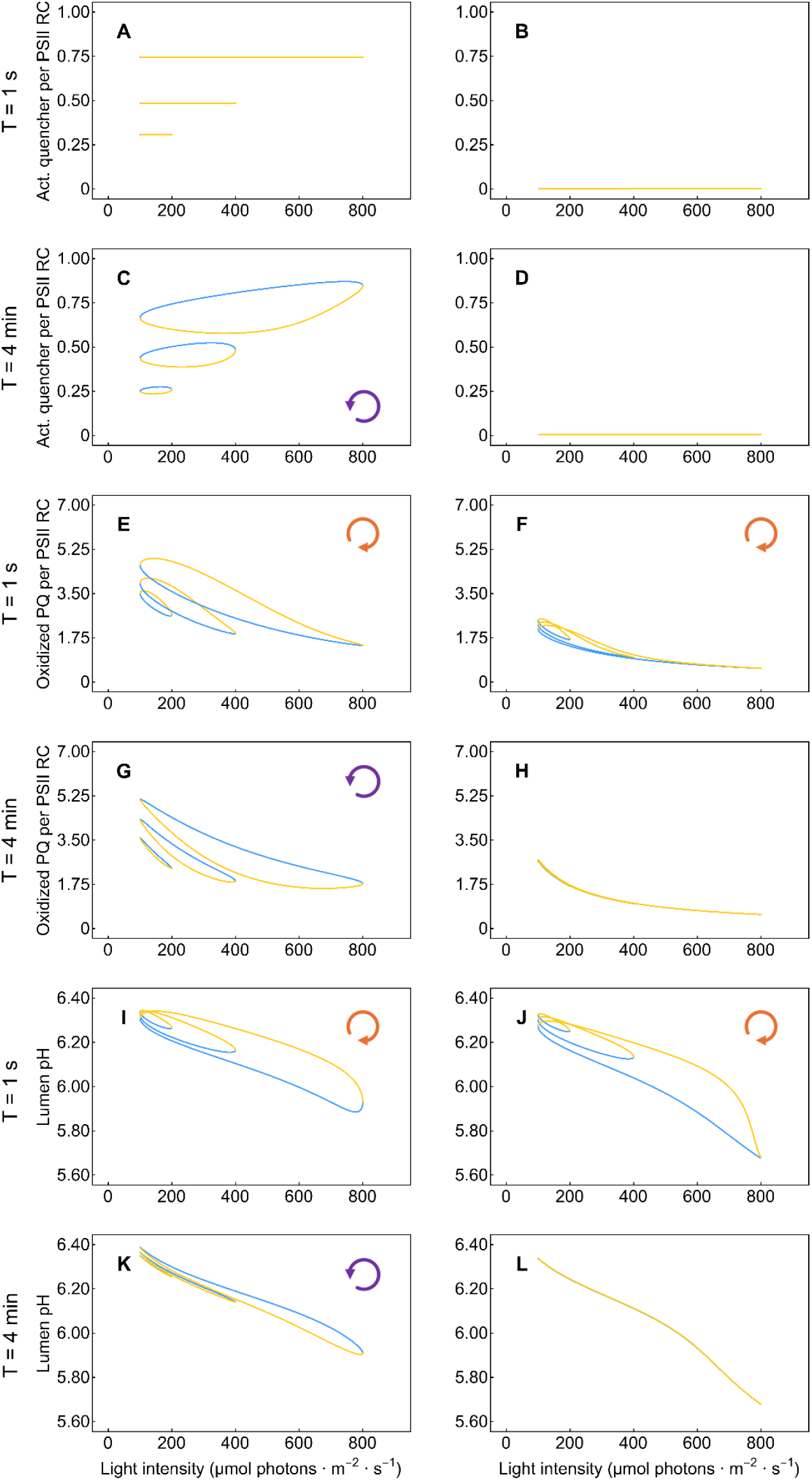
The simulated dynamics of the active quencher per PSII reaction center, the FQact(t) variable of the model (panels A-D), of oxidized PQ pool per PSII reaction center, the PQ(t) variable of the model (panels E-H), and of the lumen pH, calculated as - log_10_(H_L_(t)) of the model (panels I-L). The respective oscillation periods T = 1 s and 4 min periods are marked in the y-axis legends. The panels of the left column show simulations with qE mechanism active, while those of the right column panels depict simulations for FQact = 0. The color code of the simulated curves is the same as in the previous figures. The loop arrow at the upper left corner of each panel indicates the orientation of the ChlF(t) loops.

The simulations showed that the independent variable FQact(t) representing the active quencher did not change when the light oscillation period was short, T = 1 s (Fig. 5A). This suggests that in such rapid light oscillations, the activation and deactivation processes of qE are unable to keep pace with changes in light and respond only to the average light intensity. The inability of the qE quencher to activate and deactivate rapidly in fast light oscillations was due to the slow response of the qE quencher to the changes in the lumen pH (Fig. 5I). This explains the absence of regulatory hysteresis with T = 1 s.

The steady-state quencher variable FQact(t) was predicted to be high in the high-amplitude oscillations and low in the low-amplitude rapid oscillations (Fig. 5A), always correlating with the average light intensity during the oscillations. This explains why the WT’s ChlF(t) loops were, on average, high in the low-amplitude oscillations and low in the high-amplitude oscillations (Fig. 4A, B).

The constitutive hysteresis forming the stimulus-response ChlF(t) patterns in the short-period oscillations, T = 1 s is caused by delays and, i.e., hysteresis in reducing and oxidizing the PQ pool both in the presence (Fig. 5E) and in the absence of qE (Fig. 5F). The maximal oxidation of the PQ pool (Fig. 5E, F), and, hence, the minimal ChlF(t) (Fig. 4B, F) appears in the rapidly oscillating light, T = 1 s during the ascending light phase, i.e. the response is delayed after the stimulation, and, therefore, the stimulus-response loop marking this constitutive hysteresis has the counter-clockwise orientation. This happens in the WT (Fig. 3A) as well as in the mutants (Fig. 3B, C) and is also simulated by the model (Fig. 4B, F).

In the long-period oscillations with T = 4 min, the quencher activation represented by the variable FQact(t) was tracing the light oscillation phase (Fig. 5C). The minimum of the quencher activation was occurring, however deep in the ascending light phase that is marked in Fig. 5C by the yellow line color. This means that the quenching response was delayed after the light oscillation; the ChlF(t) maximum due to minimal quenching also appeared in the ascending light phase and, therefore, before the light maximum was reached (Fig. 4C, D). This apparent shift between ChlF(t) and light oscillation was reflected in the clockwise orientation of the regulatory hysteresis.

Unlike the delayed lumen pH responses observed under short-period oscillations (Fig. 5I), the lumen pH response in long-period oscillations was nearly synchronized with light intensity changes (Fig. 5K). However, even in the long-period oscillations, the activation and deactivation of the qE quencher (Fig. 5C) still lagged behind the changes in lumen pH (Fig. 5K).

## 4 Discussion

### 4.1 The constitutive hysteresis of ChlF(t) reflects a kinetic limitation in the electron transport chain

The dynamics of ChlF(t) in the rapidly oscillating light (T ≤ 10 s) exhibit constitutive hysteresis, characterized by a delayed ChlF(t) response to the light stimulation occurring in WT plants (Fig. 2A-C) as well as in the mutants (Fig. 3A-C). It is the filling and emptying of the primary reactant pools (Fig. 5E) (Nedbal & Koblížek, 2006; Rascher & Nedbal, 2006; Kalaji *et al*., 2012) rather than regulation (Fig. 5A) that causes the delayed response and the constitutive hysteresis.

The constitutive hysteresis is frequency-dependent, having less influence when the periods of the light oscillations are much longer (T > 10 s) than the characteristic times of filling and emptying of the reactant pools. With this, the model predicts no hysteresis in the PQ pool redox state (Fig. 5H) or in lumen pH (Fig. 5L) for long periods.

In addition to the hysteresis loop, the response is also formed by saturation of the photosynthetic reactions in the high light range. It is important to note that, although the qE regulation cannot keep pace with the rapidly oscillating light, it is responding to the average light intensity (Fig. 5A). The quencher activation is high in light that oscillates between 100 and 800 µmol photons·m^-2^·s^-1^ and decreases with the oscillation maxima dropping to 400 and 200 µmol photons·m^-2^·s^-1^ (Fig. 5A).

### 4.2 The regulatory hysteresis of ChlF(t) reflects a delay in the qE response

The qE regulation in WT plants could follow the light oscillations when the periods were T = 30 s and longer. The quenching, however, lagged the light oscillation, which led to the ChlF(t) maximum occurring before the maximum of the light and, therefore, to a change of the ChlF(t) loop orientation. The ChlF(t) loop orientation changed from counter-clockwise with the short oscillation periods (Fig. 2A-C) to clockwise with the long periods (Fig. 2D-H) because the latter response was primarily formed by the regulation lag (Fig. 5C). This is in agreement with the measurements that applied saturated flashes during the light oscillations and that revealed a delay of about 15 s during the ascending light phase of T = 1 min light oscillations (Shimakawa & Miyake, 2018; Lazár *et al*., 2022; Niu *et al*., 2024).

The oscillation period of 1 min was already long enough for extensive periodic activation and deactivation of the quenching mechanisms and, yet, still comparable to the lag in the regulatory response (Shimakawa & Miyake, 2018; Lazár *et al*., 2022; Niu *et al*., 2024). This made the regulatory hysteresis in the loop in Fig. 2E dominant. Further increasing the oscillation period to several minutes in Fig. 2F-H led, particularly in the high-light range, to the narrowing of the hysteresis loops that can be explained by a fully developed qE that can follow the slowly oscillating, strong light with a negligible delay.

The regulatory hysteresis dominated the ChlF(t) response to 1-min light oscillations not only in the WT (Fig. 3D) but also in the *npq1* mutant (Fig. 3E), both competent in the PsbS-dependent qE. The absence of a similarly strong hysteresis in the *npq4* mutant (Fig. 3F) indicates that the observed regulatory hysteresis depends on the dynamics of PsbS protein activation and deactivation. The *npq1* mutant (Fig. 3E) exhibited higher average ChlF(t) than the WT (Fig. 3D), suggesting that zeaxanthin-dependent qE, though not dominating the regulatory dynamic response, reduces the amplitude of ChlF(t) changes and suppresses oscillation in the photosynthesis system under oscillating light. However, zeaxanthin-dependent qE alone fails to induce effective dynamic regulation in the absence of PsbS protein (Fig. 3F).

Constitutive and regulatory hysteresis were observed in the WT in high-as well as in low-light oscillations (Fig. 3A, D, G). Constitutive hysteresis was observed in the *npq1* and *npq4* mutants also with all light oscillation amplitudes when the oscillation periods were short (Fig. 3B, C), confirming that the phenomenon depends on the primary reactions, not on qE regulation. Interestingly, however, the regulatory hysteresis that was apparent in the *npq1* mutant in the high- and medium-amplitude light oscillations was not expressed when the light oscillated only in the sub-saturating intensity range (Fig. 3E & Fig. SI-5 E-H). This may be tentatively interpreted by zeaxanthin’s role in modulating the relationship between qE and lumen pH (Noctor *et al*., 1991; Noctor *et al*., 1993). Zeaxanthin acts as an allosteric modulator of qE, altering its efficiency and kinetics by shifting the apparent pK of qE from 4.5 to 6.5 or a more alkaline pH (Crouchman *et al*., 2006; Johnson *et al*., 2008; Pérez-Bueno *et al*., 2008; Johnson *et al*., 2009; Johnson & Ruban, 2009). Existing zeaxanthin in WT plants enables qE activation at a higher lumen pH, which typically occurs in low-light oscillation, whereas in the *npq1* mutant, the absence of zeaxanthin necessitates a lower lumen pH to activate qE. These findings support the idea that zeaxanthin plays a regulatory role in qE response, and that the relationship between qE and ΔpH is non-linear and dynamically altered(Noctor *et al*., 1991; Niyogi *et al*., 1998; Johnson *et al*., 2008; Nilkens *et al*., 2010; Jahns & Holzwarth, 2012).

In previous paper (Niu *et al*., 2023), we proposed that in high-light oscillation with tested periods (100 – 800 µmol photons·m^-2^·s^-1^; 1 s – 8 min periods), zeaxanthin produced during the pre-illumination and high-light phases of oscillation cannot apparently decline during the relatively brief low-light phases of oscillation. However, this may not apply to the low-light oscillations studied here (100 – 200 µmol photons·m^-2^·s^-1^). The re-conversion of zeaxanthin to violaxanthin in darkness or low light depends on pre-illumination intensity, with higher intensity slowing down the re-conversion of zeaxanthin to violaxanthin by lowering the amount of zeaxanthin epoxidase through protein degradation (Jahns, 1995; Jahns & Holzwarth, 2012; Kress & Jahns, 2017). Low-light oscillations with long periods may allow the xanthophyll cycle to operate bidirectionally with ΔpH changes (Jahns, 1995), leading to dynamic changes in zeaxanthin concentration. The potential oscillations in zeaxanthin concentration can directly affect qE, which could also explain the difference of ChlF(t) dynamics observed between the WT and *npq1* mutant in low-light long-period oscillations (Fig. 3D, E, G, H). Further studies on the changes of xanthophyll composition and the proton motive force could clarify zeaxanthin’s role in qE regulation under oscillating light.

### 4.3 Quantitative differences between experimental data and the simulations

The BDM was able to correctly distinguish the constitutive and regulatory hysteresis of the ChlF(t) relative to the phase of the oscillating light. Yet, significant differences between data and simulations are also apparent in Fig. 4 and represent opportunities to improve the model.

The stoichiometries, rate constants, and other parameters of the applied BDM model, which were summarized in Tab.1 in Fuente *et al*. (2024), were largely collected from literature, reporting measurements with often different species that were acclimated to different environmental conditions. Some of the model parameters were only guessed or used as formal placeholders for future model amendments. Finding quantitative differences between the data on *A. thaliana* and model simulations is, therefore, not surprising. At least a part of these differences can be alleviated by a parameter selection that would be, as much as possible, using literature on the same species in similar experimental conditions. This may not always be possible also because the model used effective rate constants, e.g., k1, that lumped multiple processes involved in the electron transfer from PSII RC to the PQ pool.

Computer optimization (fitting) of the parameters could reduce the differences between the model simulations and experiments. It is important to note that this would require non-linear parameter optimization owing to the structure of the BDM. However, it would most likely remain a challenge to reproduce experimental data perfectly because of the reasons mentioned above.

Further sources of the quantitative differences between the model simulations and the experiment are the simplifications that were needed to minimize the model dimensionality. In addition to the lumped multiple processes, the model considers only one mechanism of NPQ which, with respect to values of related model parameters, describes the zeaxanthin-dependent qE. The linear electron transport beyond the PQ pool, photosynthesis control, cyclic electron transport, Calvin-Benson-Bassam cycle, and other processes are not included in the BDM and might be important for shaping the ChlF(t) signal under oscillating light. For example, the electric potential difference across the thylakoid membrane was reported to play a role in the oscillations of ChlF(t) under sinusoidal illumination with the period of 1 min (Lazár *et al*., 2022). These processes should also be considered in a model for explanation of further details in photosynthesis dynamics in the oscillating light. On the other hand, each of such potential model expansions should be critically evaluated with respect to potential model over-parametrization. It is wise to adhere to the parsimony principle and keep the model dimensionality at its minimum, which is required to explain the phenomena observed in the experiments (Fuente *et al*., 2024). However, the parsimony principle does not mean that the models must be simple. For example, this work showed that consideration of only one type of qE mechanism in the BDM is not enough to describe experimental data from *npq4* and *npq1* mutants in detail. The same would be true for the two types of cyclic electron transport pathways that were experimentally characterized by using oscillating illumination (Niu *et al*., 2023; Niu *et al*., 2024).

## Author contributions

S.M. formulated the experimental plan and contributed to the final formulation of the manuscript. Y.N. performed the experiments, analyzed the experimental data and described them in the initial version of the manuscript. D.F. participated in developing the model, did the computer simulations, described them in the initial version of the manuscript, and generated all figures. D.L. participated in developing the model and wrote the final manuscript version. L.N. formulated the approaches to the data analysis, designed the analysis routines, participated in developing the model, and wrote the final manuscript version.

## Data availability statement

Raw experimental data and BDM code are available upon request.

## SUPPORTING INFORMATION

### Model description and modification of the ChlF(t) expression

The scheme of the basic DREAM model (BDM; Fuente et al., 2024) used for simulations is shown in Fig. SI-7. We kept the definition of BDM as in the original work, but we modified the way the ChlF(t) is calculated to account for the minimal relative chlorophyll fluorescence yield in the light-adapted state F_0_′(t). Using the formula derived by Oxborough and Baker (1997), one can express this dependent variable as:

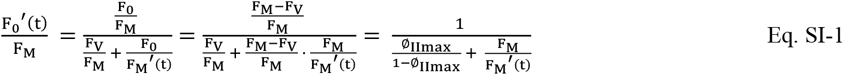

where Φ_IImax_ = F_V_/F_M_ is the maximum quantum yield for PSII photochemistry in the dark-adapted state and F_V_ and F_M_ are the parameters measured by PAM techniques. Here, we use F_V_/F_M_ = 0.832 (Björkman and Demmig, 1987).

The maximal relative chlorophyll fluorescence yield in the light-adapted state F_M_′(t) is reduced relative to the respective parameter in the dark-adapted state F_M_ by the non-photochemical quenching that is characterized in BDM by the maximum quenching level FQ_max_ and by the independent variable FQ_act_(t) that reflects the activity of the quencher during the light oscillations:

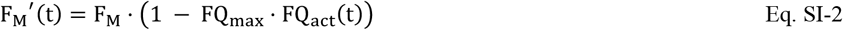

Substituting Eq. SI-2 in Eq. SI-1, one obtains:

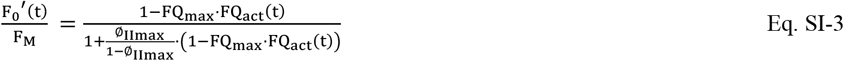

Similarly, the normalized relative variable chlorophyll fluorescence yield can be expressed as:

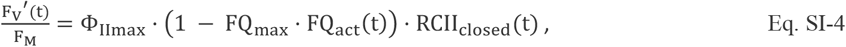

where RCII_closed_(t) is the fraction of closed (with reduced Q_A_) reaction centers of PSII at time t.

Adding 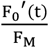 from Eq. SI-3 and 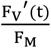 from Eq. SI-4, one obtains for total ChlF(t) Eq.2 as it is used in the main text:

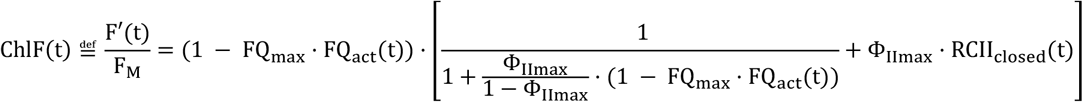

Unlike Fig.2 in the main text, the numerical fit does not interpolate the data here. Comparing this figure with Fig.2 shows that the numerical fitting process by the function from Eq.2 does not distort the ChlF(t) dynamic pattern observed in the experiments. The fitting by Eq.2 is done to compare analytical functions characterizing the experiment and the simulations.

**Figure SI-1.**
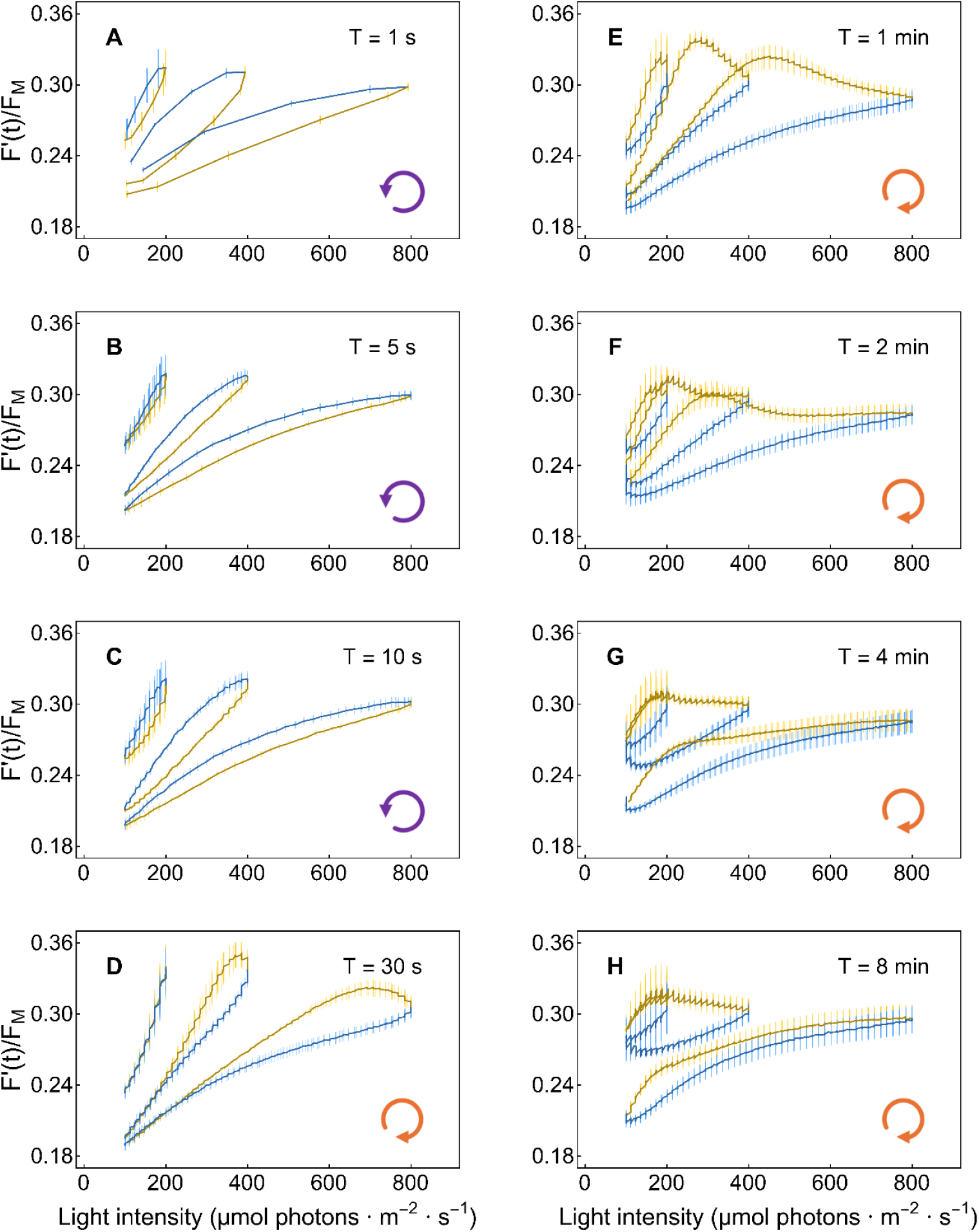
The dynamics of the original experimental data on ChlF(t) obtained with the WT *A. thaliana* Col-0. ChlF(t) is shown as a function of the intensity of PAR that oscillates with different periods and amplitudes. The dynamics represent an average of three independent biological replicates, with error bars indicating standard errors (n = 3).

**Figure SI-2.**
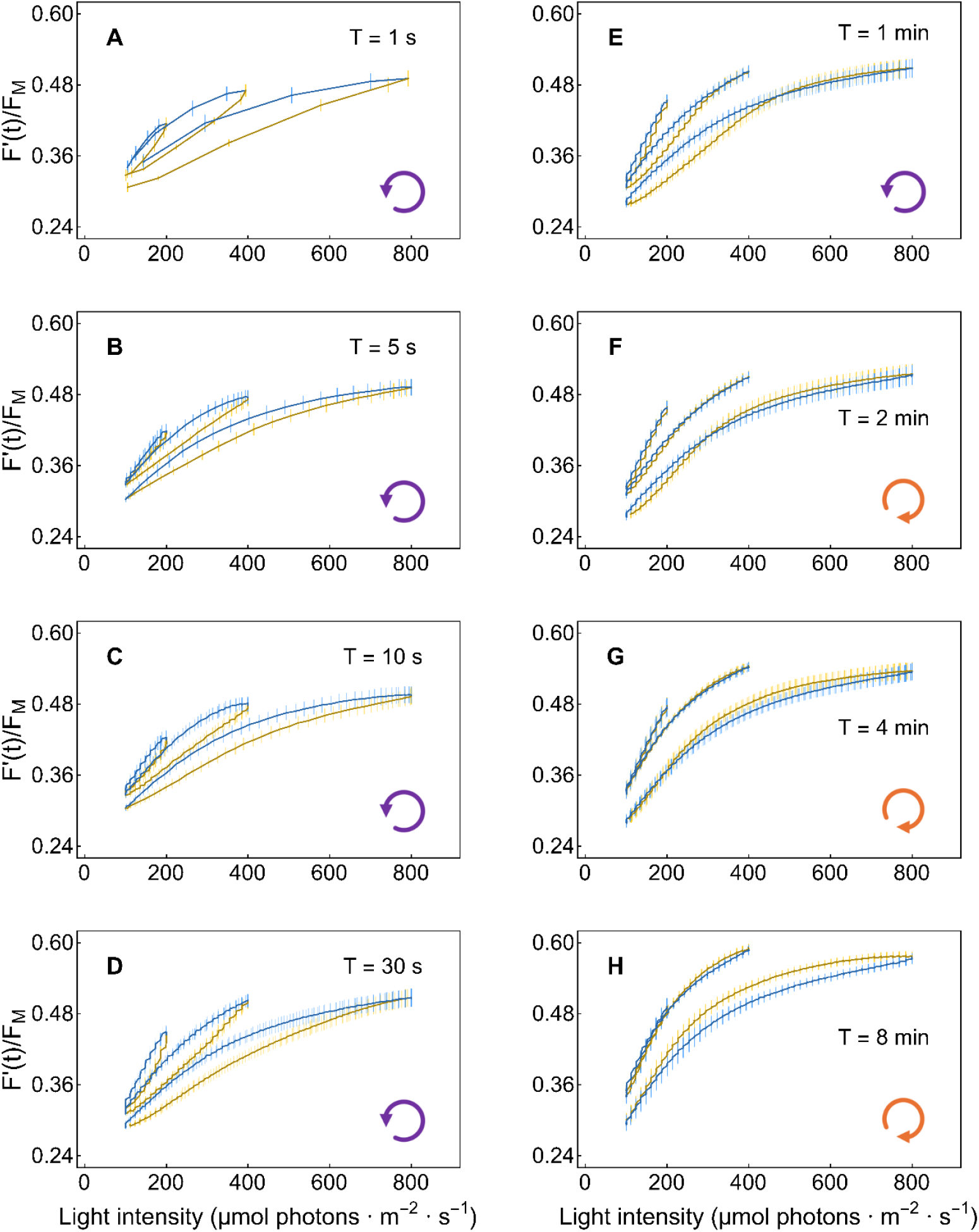
The dynamics of the original experimental data on ChlF(t) obtained with the *A. thaliana npq4* mutant. The marking and legends are the same as in Figure SI-1.

**Figure SI-3.**
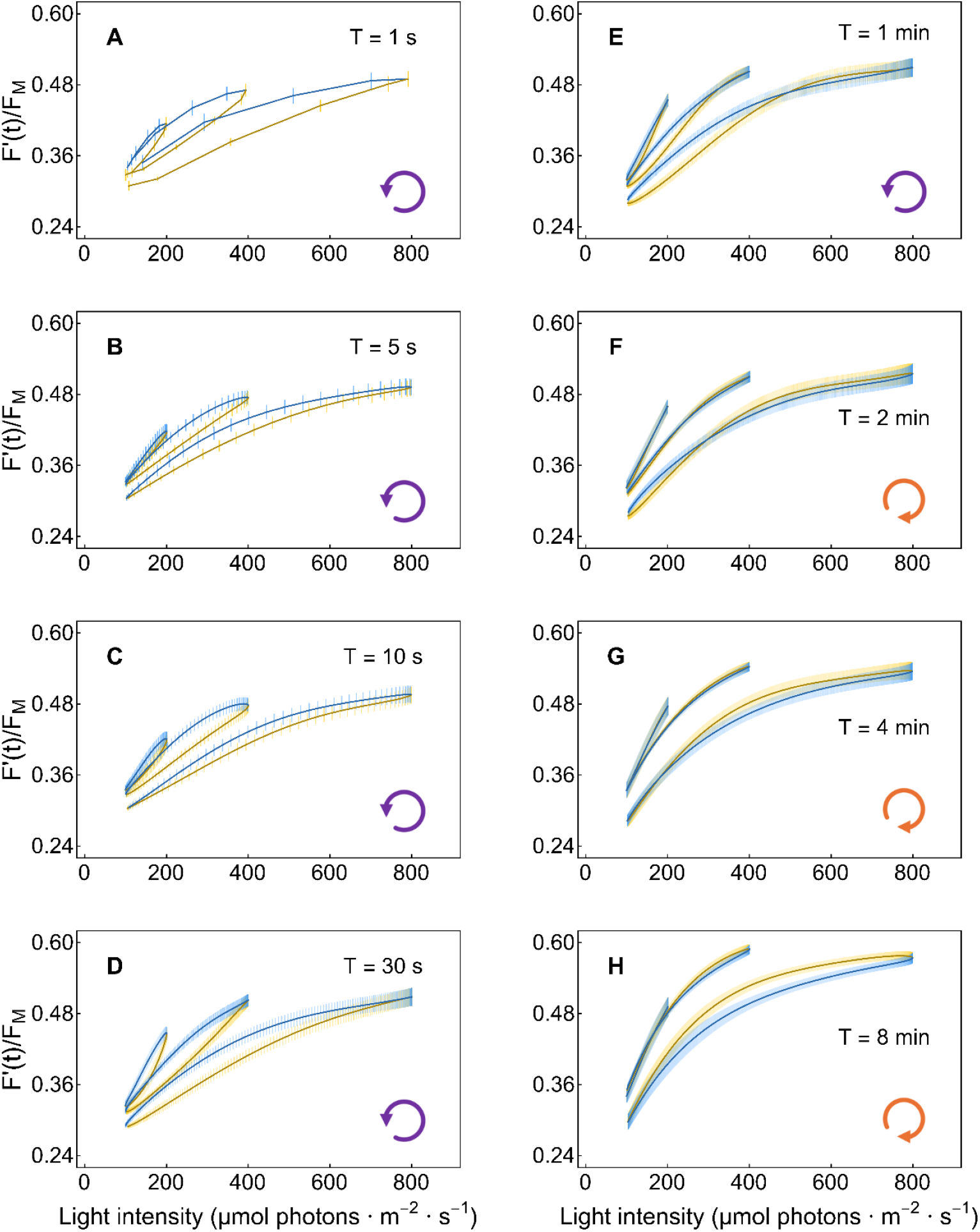
Data on *A. thaliana npq4* mutant represented in Figure SI-2 fitted by the function in Eq.1, the fits averaged and analyzed for experimental error and represented here by resulting analytical function.

The agreement between Fig. SI-2 and SI-3 shows that the numerical fitting did not change the dynamic pattern.

**Figure SI-4.**
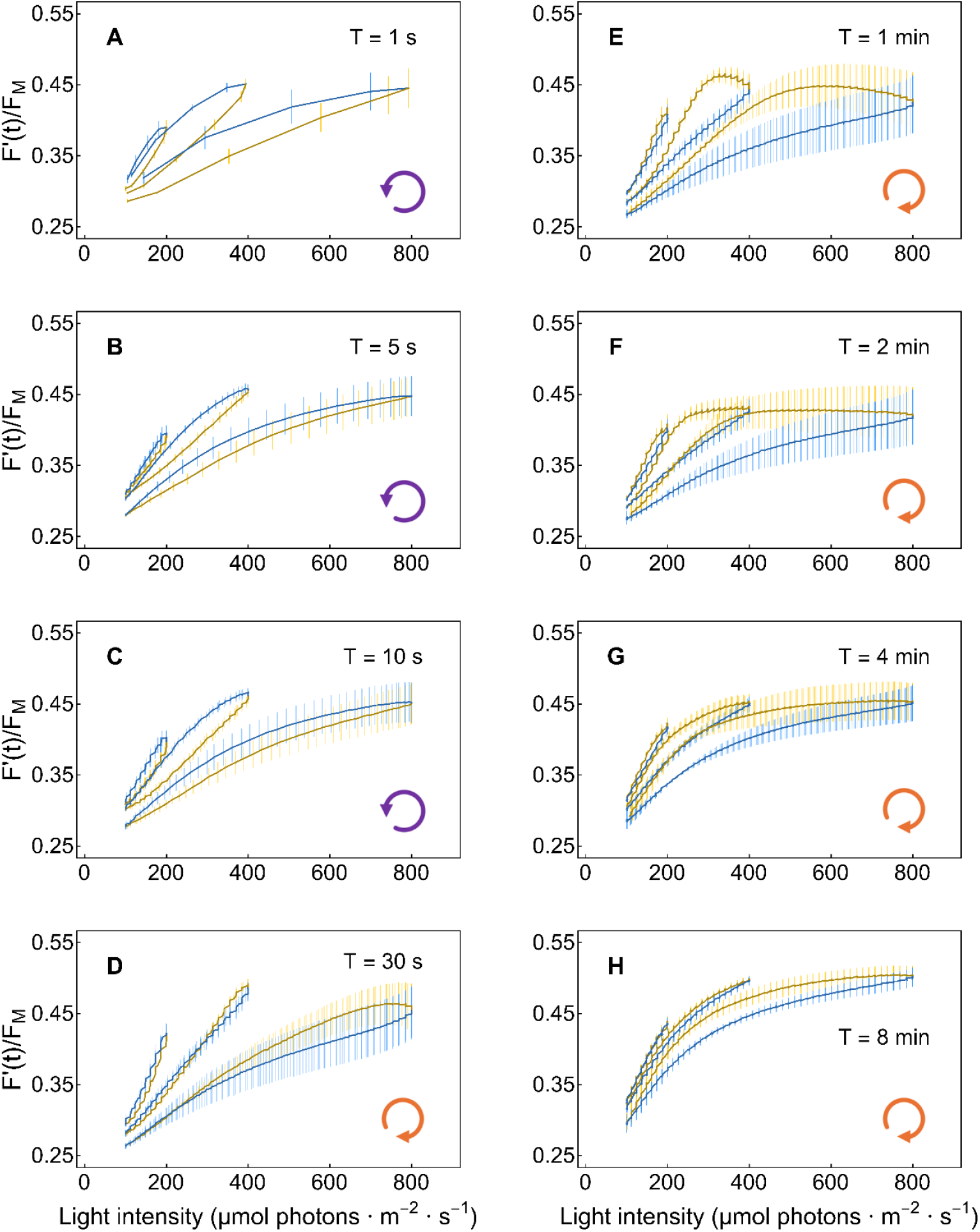
The dynamics of the original experimental data on ChlF(t) obtained with the *A. thaliana npq1* mutant. The marking and legends are the same as in Figure SI-1.

The agreement between Fig. SI-4 and SI-5 shows that the numerical fitting did not change the dynamic pattern.

**Figure SI-5.**
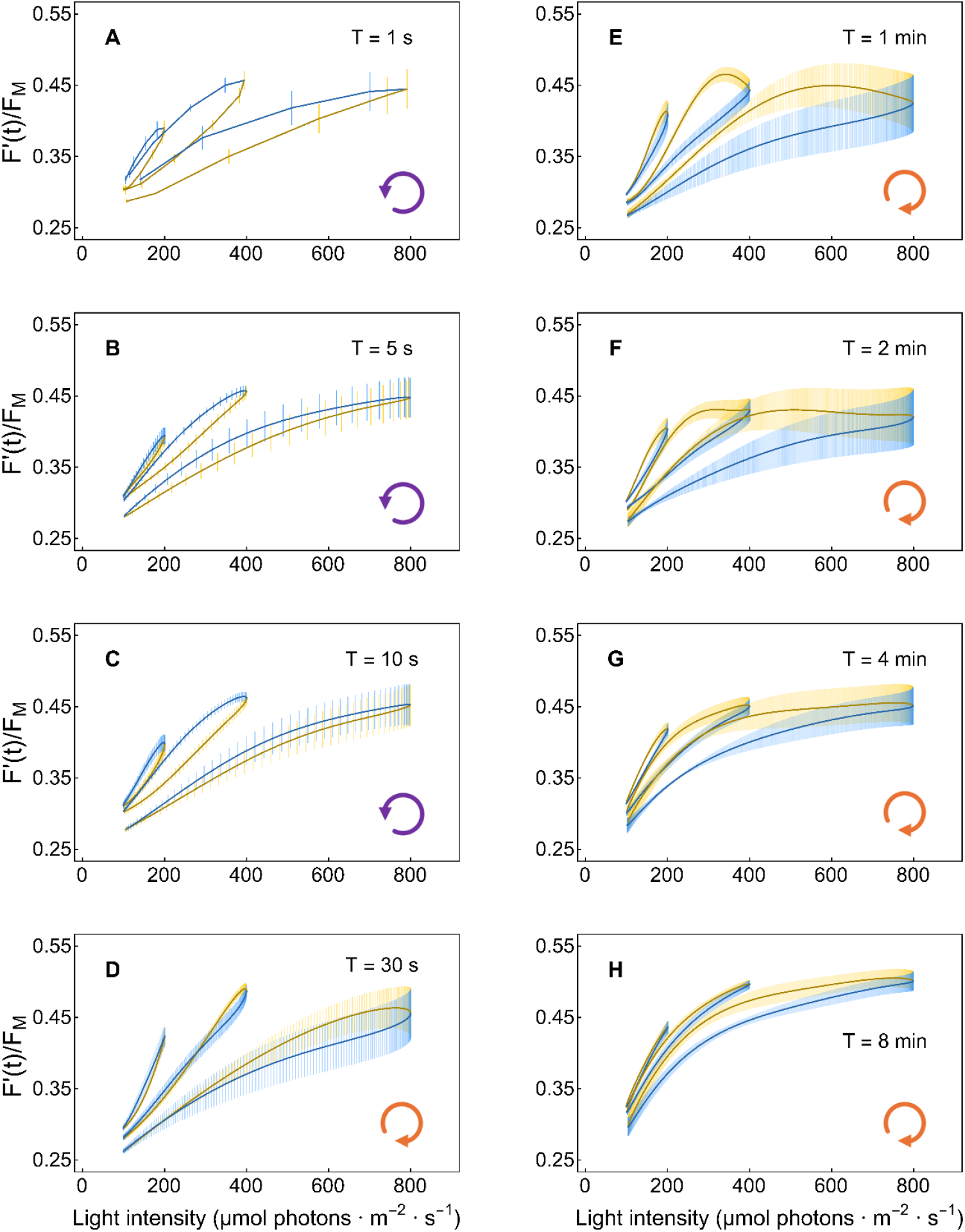
Data on *A. thaliana npq1* mutant represented in Figure SI-4 fitted by the function in Eq.1, the fits averaged and analyzed for experimental error and represented here by resulting analytical function.

**Figure SI--6.**
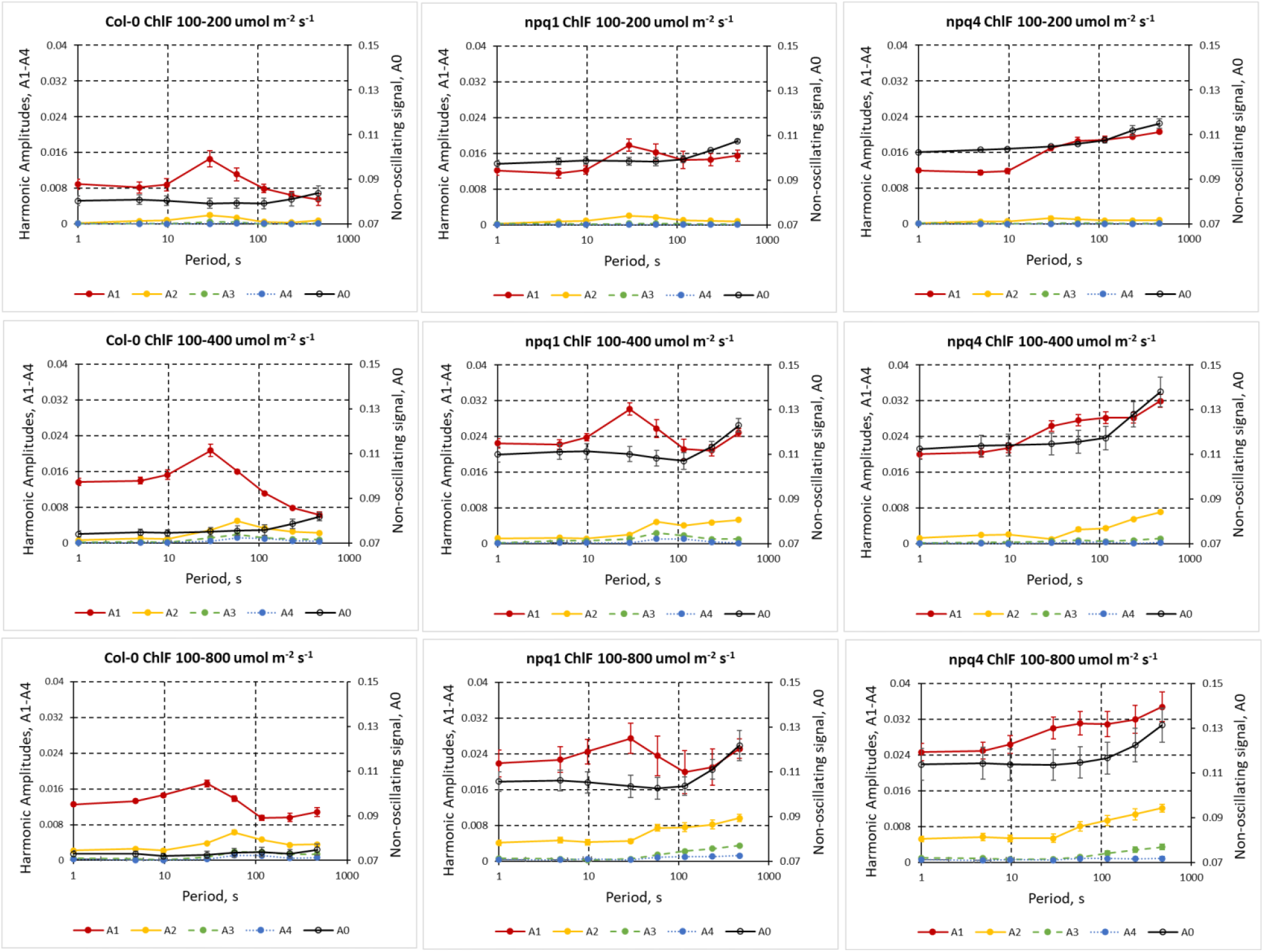
The experimentally measured ChlF(t) response to harmonic light stimulation was deconvoluted by fitting the function defined in Eq.2. The numerical fits yielded the constant component A_0_, the fundamental oscillatory amplitude A_1_, and the three upper harmonic amplitudes A_2_, A_3_, and A_4_. The individual graph panels show how the offset A_0_ (black lines) and the harmonic amplitudes A_1_ (red), A_2_ (yellow), A_3_ (green), and A_4_ (blue) changed with the period of light oscillation. The left column panels represent the WT, the middle column the *npq1* mutant, and the right column the *npq4* mutant. The amplitude of the light oscillations was low in the top row of panels, medium in the central row, and high in the bottom panel row.

**Figure SI-7.**
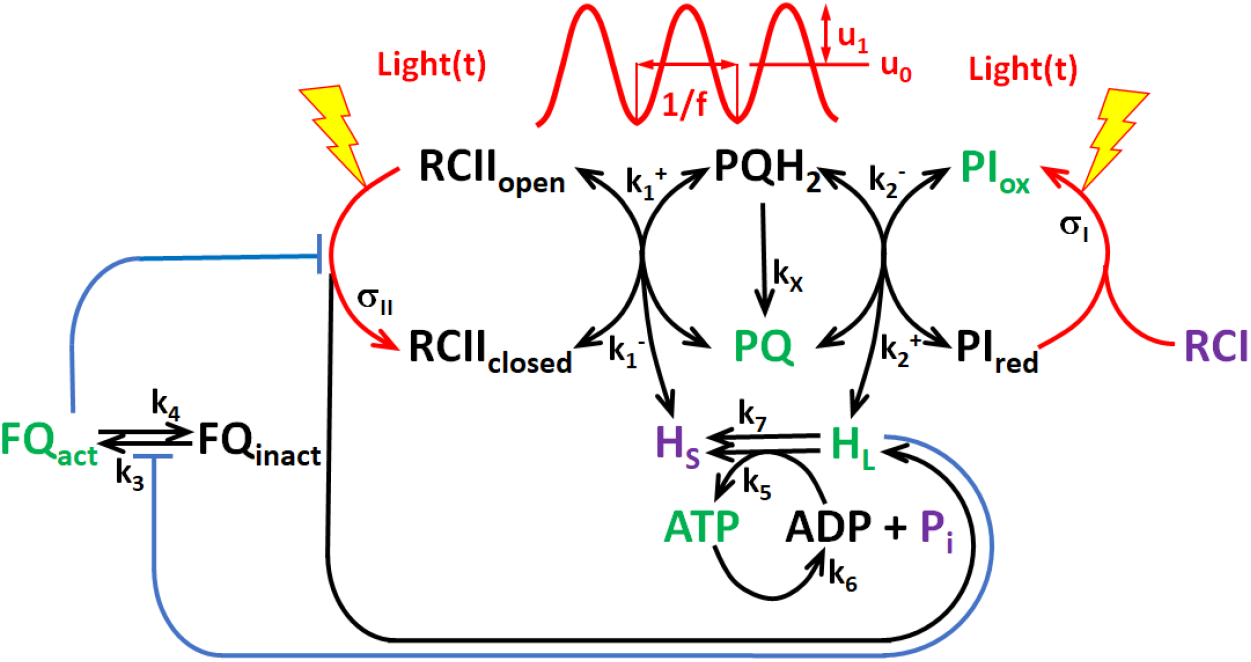
Scheme of the basic DREAM model (BDM). The red-colored sinusoidal curve represents harmonic light modulation. The oscillations of amplitude u_1_ and period 1/f (f is the frequency) are superimposed on the constant light level u_0_. The green color marks the independent variables. The black color lettering represents the dependent variables, rate and other constants. The purple color marks the model variables that are assumed to be constant in time. For details about the model see Fuente et al. (2024).

## Footnotes

1 It is sometimes also called the input-output relationship, as in Frank 2013.

2 Harmonic functions of various periods constitute a unique base in the Hilbert space of all functions that may represent various light fluctuating patterns (Schwartz 2008). They are unique by their orthogonality that might have an analogue in the orthogonality of the usual X-Y-Z Cartesian coordinates. The fact that the X-, Y-, and Z-axis are perpendicular, i.e., orthogonal means that every position in our 3D space can be characterized by 3 unique numbers (x, y, z). Similarly, any relevant function representing a light pattern can be represented by a linear combination of harmonic functions of various frequencies.

3 We use the term “steady-state dynamic pattern” to describe the response of a system that exhibits continuous motion or oscillation while maintaining a constant overall behavior. Here, it was when A_0_, A_1,_ A_2,_ A_3,_ A_4_ and τ_1_/T, τ_2_/T, τ_3_/T, τ_4_/T were not changing over time or when the changes were small and smoothed out by averaging over several oscillation periods.

